# Fast 3D printing of large-scale biocompatible hydrogel models

**DOI:** 10.1101/2020.10.22.345660

**Authors:** Nanditha Anandakrishnan, Hang Ye, Zipeng Guo, Zhaowei Chen, Kyle I. Mentkowski, Jennifer K. Lang, Nika Rajabian, Stelios T. Andreadis, Zhen Ma, Joseph A. Spernyak, Jonathan F. Lovell, Depeng Wang, Jun Xia, Chi Zhou, Ruogang Zhao

**Affiliations:** Department of Biomedical Engineering, University at Buffalo, The State University of New York, Buffalo, NY 14260 USA; Department of Industrial and Systems Engineering, University at Buffalo, The State University of New York, Buffalo, NY 14260 USA; Department of Medicine, Division of Cardiology, Jacobs School of Medicine and Biomedical Sciences, University at Buffalo, Buffalo, NY, 14203 USA; VA WNY Healthcare System, Buffalo NY, 14215 USA; Department of Chemical and Biological Engineering, University at Buffalo, The State University of New York, Buffalo, NY 14260 USA; Department of Biomedical and Chemical Engineering, Syracuse Biomaterials Institute, Syracuse University, Syracuse, NY 13244 USA; Department of Cell Stress Biology, Roswell Park Comprehensive Cancer Center, Buffalo, NY USA

**Keywords:** 3D bioprinting, stereolithography, engineered tissues, hydrogels, rapid printing, large-sized model

## Abstract

Large scale cell-laden hydrogel models hold great promise for tissue repair and organ transplantation, but their fabrication is faced with challenges in achieving clinically-relevant size and hierarchical structures. 3D bioprinting is an emerging technology, but its application in large, solid hydrogel fabrication has been limited by the slow printing speed that can affect the part quality and the biological activity of the encapsulated cells. Here we present a Fast hydrogeL prOjection stereolithogrAphy Technology (FLOAT) that allows the creation of a centimeter-sized, multiscale solid hydrogel model within minutes. Through precisely controlling the photopolymerization condition, we established low suction force-driven, high-velocity flow of the hydrogel prepolymer that supports the continuous replenishment of the prepolymer solution below the curing part and the nonstop part growth. We showed that this process is unique to the hydrogel prepolymer without externally supplemented oxygen. The rapid printing of centimeter-sized hydrogel models using FLOAT was shown to significantly reduce the part deformation and cellular injury caused by the prolonged exposure to the environmental stresses in layer-by-layer based printing methods. Media perfusion in the printed vessel network was shown to promote cell survival and metabolic function in the deep core of the large-sized hydrogel model over long term. The FLOAT is compatible with multiple photocurable hydrogel materials and the printed scaffold supports the endothelialization of prefabricated vascular channels. Together, these studies demonstrate a rapid 3D hydrogel printing method and highlight the potential of this method in the fabrication of large-sized engineered tissue models.

## Introduction

Large scale cell-laden hydrogel models hold great promise for tissue repair and organ transplantation, but their fabrication is faced with challenges in achieving clinically-relevant size and hierarchical structures (1). 3D bioprinting is an emerging technology for hydrogel fabrication and has been successfully used to create hydrogel models with biomimetic structures and functions (2, 3); however, its application in large, solid hydrogel fabrication has been limited by the slow printing speed that can affect the part quality and the biological activity of the encapsulated cells (4, 5).

Due to the point-by-point deposition process used in nozzle-based bioprinting techniques, extended printing time is required to fabricate a large-sized model with fine structures (6, 7). Prolonged exposure of the encapsulated cells to a variety of printing-induced environmental factors, such as the shear stress, the low oxygen level and the temperature shock, has been shown to cause serious cellular injury and cell death (8, 9). The effort to improve the printing resolution by using small diameter nozzles can cause further damage to the cells (9, 10). Additionally, due to the low mechanical strength of the hydrogel scaffold materials, it is very challenging for point-by-point deposition methods to create overhanging or hollow structures such as vascular channels inside solid parts. To address this limitation, Atala group utilized rigid polymeric scaffolds to support the printing of cell-laden hydrogel materials (11), and Feinberg group extruded hydrogel material in a secondary supporting hydrogel to print biomimetic structures such as a heart chamber (12); however, these approaches suffer from either the high rigidity of the supporting material or the complexity of the post-processing steps. Although extrusion printing of dissolvable templates composed of sacrificial materials such as fugitive inks and carbohydrate glass has enabled the creation of perfusable vascular channels in casted hydrogel constructs (13–15), this approach has very limited capacity to create fine tissue structures other than vascular channels due to the simple casting method used.

Digital mask projection-stereolithography (MP-SLA) is a photopolymerization-based, layer-by-layer 3D printing technology that features multi-scale fabrication capacity with high spatial resolution, allowing the bulk geometry and fine structure of a complex 3D model to be built through one single process (16, 17). The liquid resin provides natural self-support for the fabrication of hollow structures. This approach has been used to fabricate hydrogel models such as nerve conduits and muscle-powered biobots. Recently, multivascular networks have been created in hydrogels by controlling the spatial resolution of hydrogel photopolymerization using selected food dye photoabsorbers (18–20); however, the layer-by-layer process used in these studies limited the printing speed, which can potentially cause dehydration-induced part deformation and reduced cell viability during the fabrication of large-sized hydrogel parts. Recently, the development of continuous liquid interface production (CLIP) technology drastically increased the fabrication speed of MP-SLA through continuously building the layers of a 3D part immediately above a “dead zone” formed by oxygen inhibition of photopolymerization (21). In the dead zone, the flow of liquid water-insoluble-resin (WI-resin) enables continuous material replenishment at the polymerization interface. However, due to the low fluidity of the WI-resin material and the corresponding large suction force at the curing interface, the fabrication ability of the CLIP technology is limited to thin-walled parts (21–23). The fabrication of a centimeter-sized solid hydrogel part has not yet been achieved using CLIP.

In this work, we established low suction force-driven, high-velocity flow of the hydrogel prepolymer for continuous MP-SLA printing through precisely controlling the photopolymerization condition. The high-velocity flow supports the continuous replenishment of the prepolymer solution below the curing part and the nonstop part growth. This method, the Fast hydrogeL prOjection stereolithogrAphy Technology (FLOAT), allows the creation of a centimeter-sized, multiscale solid hydrogel model within several minutes. We showed that this process is unique to the hydrogel prepolymer solution and cannot be achieved using resin without externally supplemented oxygen. The rapid printing of centimeter-sized hydrogel models using FLOAT was shown to significantly reduce the part deformation and cellular injury caused by the prolonged exposure to the environmental stresses in layer-by-layer based printing methods. Media perfusion in the printed vessel network was shown to promote cell survival and metabolic function in the deep core of the large-sized hydrogel model over long term. The FLOAT is compatible with multiple photocurable hydrogel materials and the printed scaffold supports the endothelialization of prefabricated vascular channels.Together, these studies demonstrate a rapid hydrogel 3D printing method and highlight the potential of this method in the fabrication of large-sized engineered tissue models.

## Results

### Control the photopolymerization conditions to enable low suction force-driven, high-velocity flow of the hydrogel prepolymer

The hydrogel photopolymerization is affected by a number of parameters including exposure energy, printing speed, pre-polymer concentration, photoinitiator absorption coefficient and photo-absorber coefficient. We studied their effects and carefully adjusted the relative ratios of these parameters to formulate the optimal photopolymerization condition for desired printing performance. We first studied the effect of different photopolymerization conditions on the flow of the hydrogel prepolymer solution. Three different curing conditions were developed by varying the photoinitiator (Lithium phenyl-2,4,6-trimethylbenzoylphosphinate, LAP) concentration while keeping the photoabsorber concentration and exposure energy constant. The motion of the fluorescence microbeads suspended in the prepolymer solution was tracked to understand the flow dynamics during printing. We showed the formation of a high-velocity directional flow of the prepolymer solution in an uncured liquid layer beneath the curing part with 0.6% LAP (Figure 1A, B, Movie S1,2). This high-velocity flow supports the continuous replenishment of the prepolymer solution below the curing part and the nonstop part growth. Measured liquid flow velocity is in the order of several hundred micrometers per second and is affected by the upward motion speed of the part and the concentration (viscosity) of the prepolymer solution (Figure 1C, Fig. S1B). The highest flow velocity was observed in low concentration solution (20% PEGDA) under high printing speed (125 μm/s), while the lowest flow velocity was observed in high concentration solution (80% PEGDA) under low printing speed (50 μm/s). Regardless of the printing speed and liquid concentration, the flow velocity was found to decrease with increased z height from the glass substrate, likely because of the increased viscosity of the precursor fluid in the top region of the liquid layer where the photopolymerization occurs. The thickness of the uncured liquid layer was found to be around 600 – 800 μm (Figure 1 B, C). The fluid flow behavior is drastically altered under other photoinitiator concentrations. With 0.1% LAP, only low-velocity and random flow was observed and the part did not form properly (Fig. S1A), suggesting an under-cured condition. With 1% LAP, overly rapid curing caused the part bottom layer to stick to the glass window, thus blocking the fluid flow (Figure S1C). In this over-cured condition, later sudden detachment of the part from the glass window due to the upward motion caused the part delamination (Movie S3). The comparison between the three photopolymerization conditions shows that 0.6% LAP is optimal for the formation of continuous prepolymer flow and nonstop part growth, and thus we chose this condition for the FLOAT printing.

**Figure 1.**
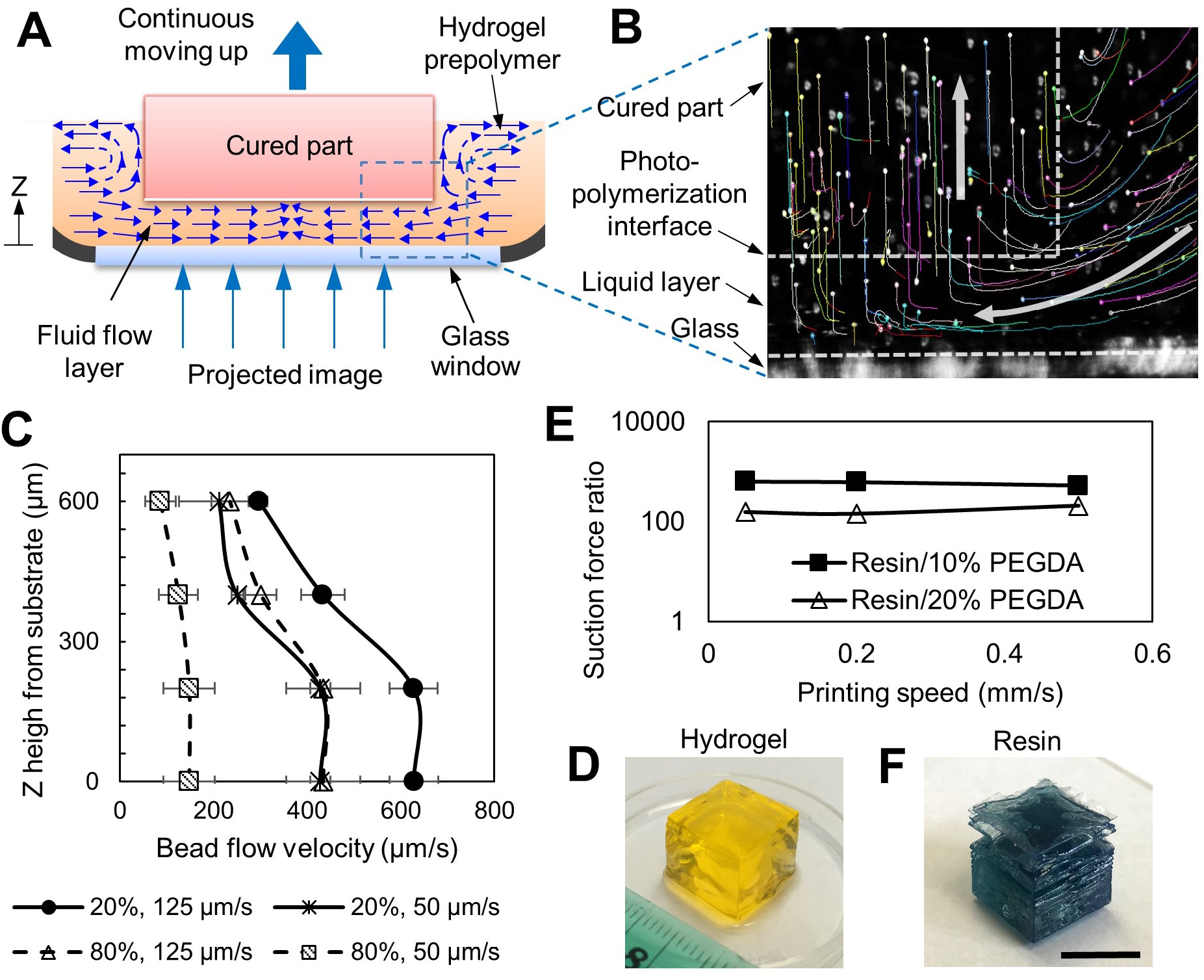
FLOAT printing is enabled by high-velocity fluid flow and low fluid suction force. (A) Schematic of the printing interface in FLOAT. Continuous replenishment of prepolymer solution below the curing part supports nonstop part growth. (B) Tracked trajectories of fluorescence microbeads during printing. White arrows show the direction of microbead motion. Microbeads were carried by the flow into a liquid layer, trapped in the hydrogel upon photocuring and then carried upwards by the cured part. (C) The flow velocity profile of 20% and 80% PEGDA prepolymer solution in uncured liquid layer under 50 and 125 μm/s printing speed. (D) A FLOAT-printed 20% PEGDA 4kDa hydrogel cube. (E) Experimentally measured suction force ratios (WI-resin / PEGDA) at various printing speeds. (F) A resin cube printed using continuous MP-SLA. Note the smooth surface and sharp edges of the hydrogel cube comparing to the rough surface and delaminated layers of the resin cube. Scale bar is 1 cm.

To understand the mechanical mechanism of the prepolymer flow, we experimentally measured the suction force at the printing interface. The suction force generated during continuous printing of a hydrogel cube under optimal curing condition was measured (Fig. S2A). The suction forces were steady and remained at a low level between 10 mN and 38 mN for 10% PEGDA part and between 41 mN and 96 mN for 20% PEGDA part under varied printing speeds (Fig. S2B, C). The steady printing process led to good part quality, as demonstrated by the printed 1 cm hydrogel cube that is featured by the smooth surface and sharp edges (Figure 1D, Fig. S2D, E). To show that this process is unique to the hydrogel material, we printed a resin cube using the same experimental setup. The suction force was unsteady and fluctuated significantly with the peak suction forces in the range of 6 N to 20 N (Fig. S3), which were several hundred times higher than those measured in hydrogel printing (Figure 1E). Highly disturbed printing process significantly impaired the quality of the resin part, causing layer delamination and part breakage (Figure 1F). Together, these suggest that the low fluid suction force in FLOAT printing helped to maintain a steady printing process, thus ensuring good part quality.

### Multiscale printing by the FLOAT

Since the photoabsorber has been shown to be critical to the printing resolution in the previous studies (25, 26), we explored the photoabsorbers that can be compatible with the FLOAT printing process. We compared commonly used photoabsorbers and studied their effects on the curing depth, which serves as a measurement of the vertical resolution. We showed that Quinoline Yellow (QY) produced the smallest curing layer thickness of 40-150 μm, thus offering the best control of the vertical resolution, due to its maximum absorption at 412 nm which is very close to the 405nm wavelength of the curing light source (Figure. 2 A,B). Since the maximum absorptions of widely used 4-hydroxy benzophenone (HMBS) and benzotriazole are between 365 and 385 nm, they were not effective in controlling the curing layer thickness. Although Tinuvin Carboprotect has a maximum absorption close to 400 nm, its poor solubility in aqueous solution reduced its efficiency in FLOAT process (Figure 2A). Increasing photoabsorber concentration was shown to reduce curing layer thickness (Figure 2B). The use of QY and other photoabsorbers such as Orange G in an optimal range was not found to significantly alter the high-velocity prepolymer flow and the continuous printing process of the FLOAT.

**Figure 2.**
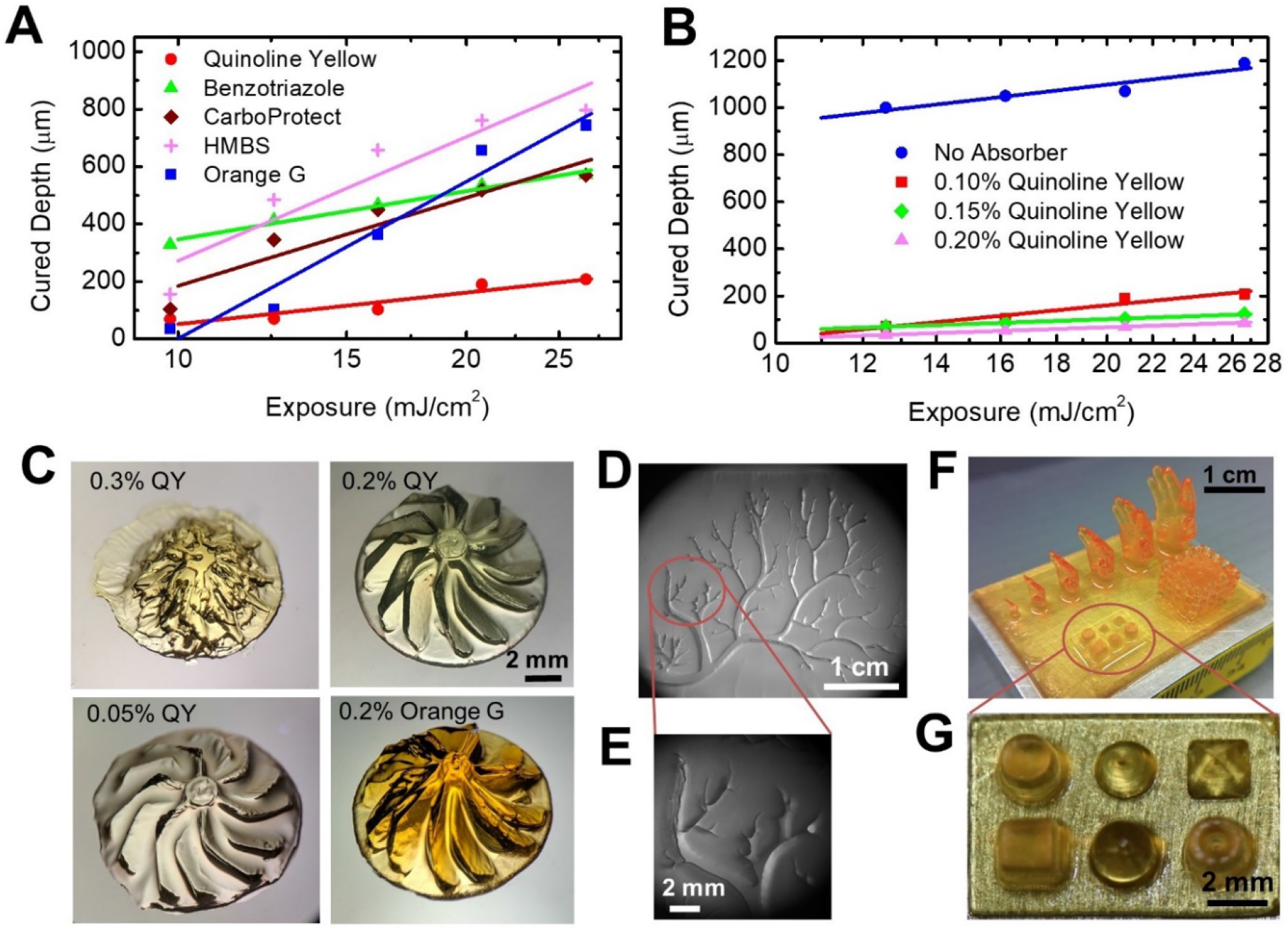
Multiscale printing by FLOAT. (A) Measured curing depths with different types of photoabsorbers. The smaller the curing depth, the greater the vertical resolution of the printed part. (B) Increasing photoabsorber concentration reduces curing depth. (C) Printing resolution demonstrated by a turbine rotor model printed under different types and concentrations of photoabsorbers. 0.3% and 0.05% QY correspond to undercured and overcured conditions respectively. 0.2% QY represents optimal curing, which results in better part quality than 0.2% Orange G. (D) FLOAT-printed vascular tree structure in a PEGDA slab with an area of 3.5 cm x 2.5 cm. (E) Enlarged view shows micrometer-scale fine structure of the vascular tree branches. (F) FLOAT-printed human hand models of different sizes, a truss model and primitives on a 3.5 cm x 2.5 cm PEGDA slab. (G) Enlarged view of the primitives. All models in this figure were printed using 20% PEGDA 4kDa.

The printing resolution and part quality of the FLOAT method were validated through printing a series of models with different geometries. We printed a turbine rotor model with an overall diameter of 1cm and individual blade thickness of 650 μm (Figure 2C, Fig. S4A-C). Since the turbine rotor blades are overhang structures with a relatively large curvature, the geometric fidelity of the blade is used as a verification of the printing quality. Undercured condition using 0.3% QY and overcured conditions using 0.05% QY were shown to lead to poor part quality as compared to the good part printed using 0.2% QY. Orange G at 0.2% allowed the printing of the overall geometry of the turbine model, but the blades are less sharp and smooth as compared to those printed with 0.2% QY (Figure 2C). HMBS is not effective in controlling the printing resolution, resulting in a heavily overcured part (Figure S4C). Figure 2D shows a 3 cm x 2.5 cm hydrogel slab containing a printed vascular tree structure with numerous branches of different widths. The widths of the largest and smallest branches are 1760 μm and 50 μm respectively (Figure 2E). A variety of 3D models were printed on a 3.5 cm x 2.5 cm rectangular hydrogel substrate that include an array of different-sized hands, a truss and an array of miniature primitives (Figure 2F, Fig. S4D). The smallest hand is 4.5 mm high with little finger diameter of 150 μm and largest hand is 17 mm high. The truss has 1 cm edge length and contains 5 struts equally distributed on each side. The primitives (cube, cone, pyramid, cylinder and dome) are about 2 mm high (Figure 2G, Figure S4E).

### The effects of rapid printing on the part quality and cellular functions in large-sized models

Here, we tested the effect of the rapid printing process of the FLOAT on the part quality and cellular health in large-sized hydrogel models. We first tested the ability of the FLOAT to rapidly fabricate centimeter-sized hydrogel models. A several centimeter-sized (2.6 x 1.7 x 5.6 cm) hydrogel human hand model was FLOAT-printed very rapidly in 20 minutes (Figure 3A-D, Movie S4). This hand model is very compliant, as demonstrated by the bending of the fingers under compression and the free recovery upon release (Figure 3E, Figure S5, Movie S5). In contrast, it took 2 hours and 6.5 hours to print the same model using the traditional layer-by-layer SLA process at 150 μm and 50 μm layer thickness, respectively. During this process, hydrogel dehydration occurred, causing distortion and detachment of the model (Figure 3F, Figure S6). The material composition and geometrical features such as the vascular channel structure of this hand model is compatible with magnetic resonance imaging (MRI), as demonstrated by the 3D reconstructed MRI scan image showing clearly the vascular channels (Figure 3G).

**Figure 3.**
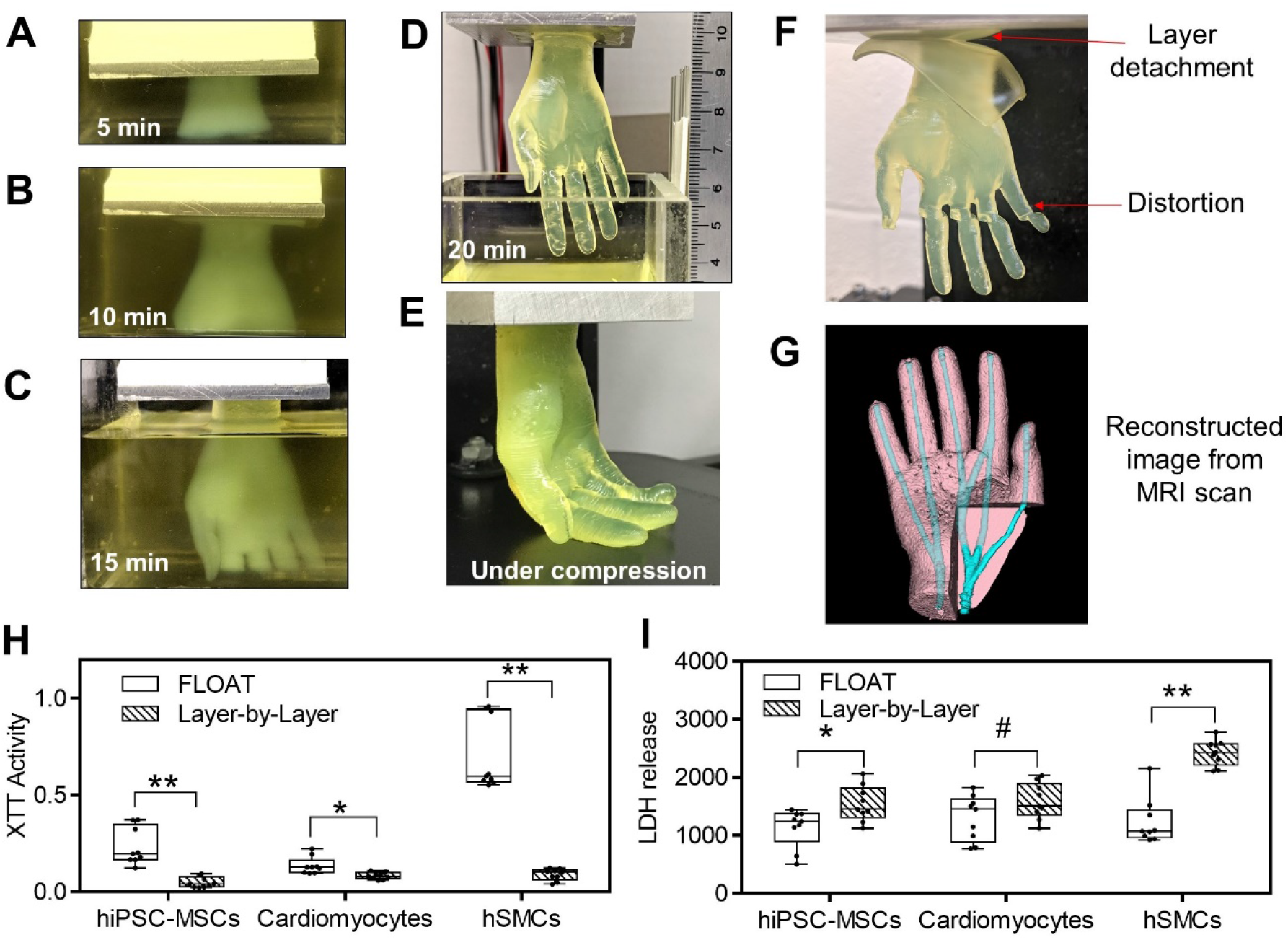
The effects of rapid printing on part quality and cellular function in large-sized models. (A-E) Demonstration of FLOAT printing process of a centimeter-sized human hand model. Sequential images of the hand model that was continuously formed in 10% PEGDA 400 Da preploymer pool at 5 min, 10 min and 15 min and completed at 20 min. It would take 6.5 hours to print the same model using traditional layer-by-layer printing method. In panel D, the length of the completed hand model is 5.6 cm. (E) Fingers were easily bent under compression, showing the compliance of the hydrogel hand model. (F) Hand model printed in 2 hours using the traditional layer-by-layer SLA process. Severe layer detachment and finger distortion occurred due to dehydration. (G) 3D reconstructed image of FLOAT-printed hand model from MRI scanning. The channels were visualized with the aid of a contrast agent. Metabolic activity and cytotoxicity measured by XTT assay (H) and LDH assay (I) shortly after the printing of centimeter-sized, cell-laden samples. Cells in FLOAT-printed samples experienced much less printing-induced injury and have much higher metabolic activity than those in layer-by-layer printed samples. The above cell-laden samples were printed using 7% GelMA plus 2% PEGDA 8k Da. n = 9. All box plots with whiskers represent the data distribution based on five number summary (maximum, third quartile, median, first quartile, minimum). **, p < 0.001; *, p < 0.05; #, p = 0.1 determined by two-tailed t-test.

We then tested the health, metabolism and viability of a variety of cell types in FLOAT-printed centimeter-sized hydrogel models (6 minutes print time), and compared the results with same-sized, cell-laden hydrogel model printed using layer-by-layer SLA (2 hours print time). Cell types that are often used in tissue engineering including human induced pluripotent stem cell-derived mesenchymal stem cells (hiPSC-MSCs), neonatal mouse cardiomyocytes and primary human skeletal muscle cells (hSMCs) were used in the tests. We showed that the averaged metabolic activities in FLOAT printed model are two to ten times higher than those in conventional layer-by-layer SLA printed model for all three cell types, as measured by the XTT assay (Figure 3H). The averaged cytotoxicity in FLOAT printed model is two to six times lower than that in conventional layer-by-layer printed model, as measured by lactate dehydrogenase (LDH) assay (Figure 3I). Together, these data suggest that cells in FLOAT-printed parts experienced much less printing-induced injury and have much higher metabolic activity than those in layer-by-layer SLA printed parts. The abilities of the FLOAT to print with uniform cell distribution and high cell density were also validated, as demonstrated in the Figure S7 and S8. The high cell density part was printed using 50 million cells per mL.

### Media perfusion in printed channels helps to maintain long-term cell functions in large-sized models

We tested the long-term cell survival and functions in FLOAT-printed, centimeter-sized tissue models. A liver-shaped model (3.5 cm x 2.5 cm x 1.5 cm) containing perfusable channels was FLOAT-printed in 5 mins using cell-laden PEGDA prepolymer conjugated with RGD peptide (Movie S6). The size of the liver model is larger than that of the rat and is comparable to the median liver size of a 26 weeks fetus (27). Visual inspection of the liver model shows smooth surface and monolithic, translucent hydrogel body without clump, bubble or delamination (Figure 4A). Perfusable channel network was fabricated in the liver model to improve the mass transport in the bulk hydrogel body. Through judiciously limiting the light exposure in designed channel areas, we fabricated the bulk body of the liver model and perfusable channels together through a single FLOAT printing process. To examine the inter-connectivity and printing fidelity of the channel network, a small molecule dye (Rhodamine B) was injected into the channels and allowed to flow through the network (Figure 4B-C, Movie S7). The cross-sectional view of the liver model shows open lumen of the channels with a height of 3 mm and width of 800 μm, which is consistent with the design value and show no sign of overcuring (Figure S9A-B). Immediately after injection, the channels contained the dye solution well and there was no dye leakage into the hydrogel interstitial space (Figure S9C). Later, time-dependent dye diffusion into the interstitial space was observed. Full diffusion of the dye throughout the entire liver model was achieved over a 30 min period (Figure S9D), as demonstrated by the gradual equilibration of the fluorescent dye intensity between the vascular channels and adjacent interstitial spaces during this period (Figure S9E).

**Figure 4.**
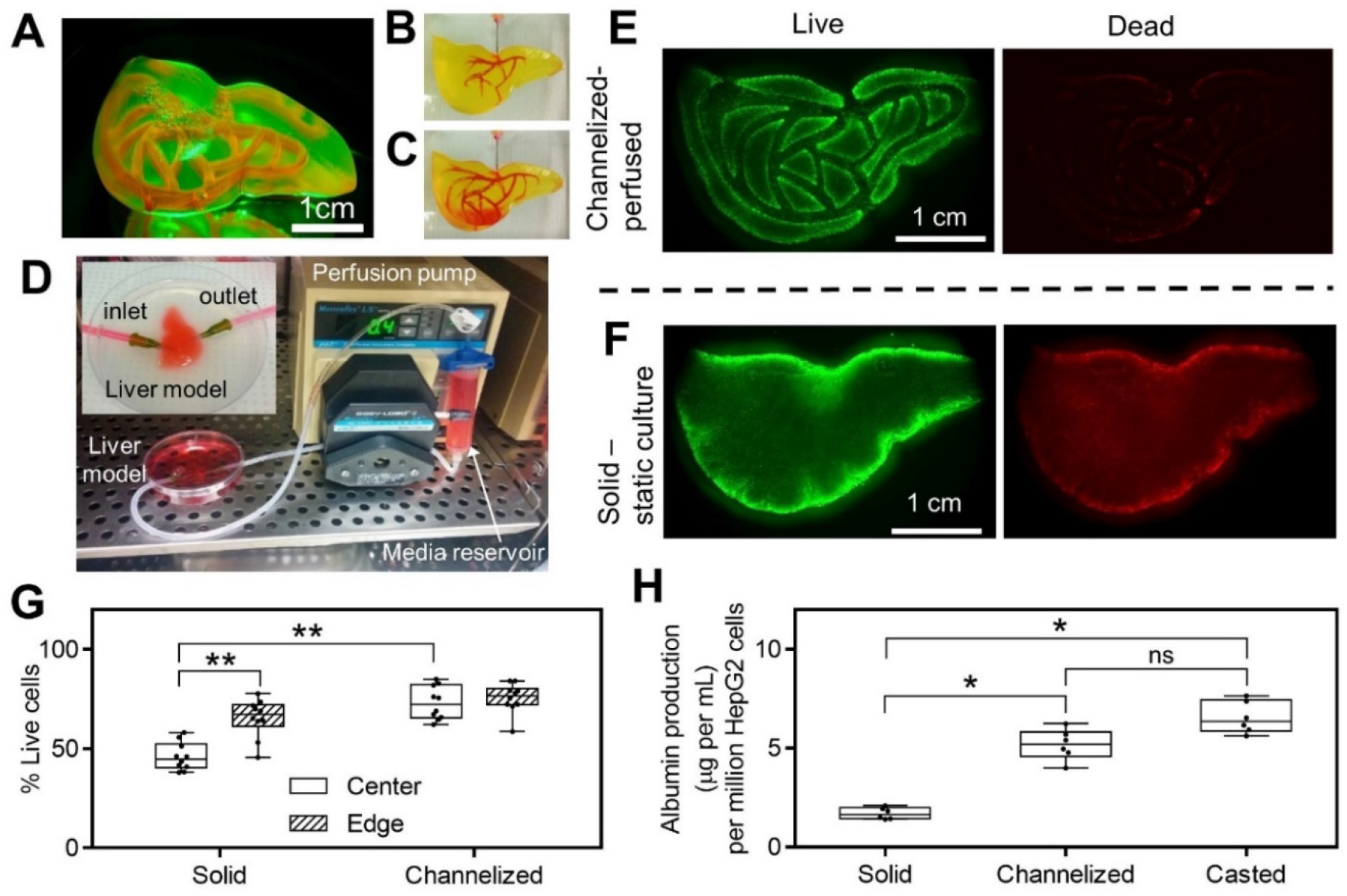
Media perfusion in printed channels helps to maintain long-term cell functions. (A) A liver model with smooth surface and monolithic, translucent hydrogel body was printed using 15% PEGDA 4 kDa. Vascular channel network was filled with Rhodamine B and visualized under fluorescence. At the beginning (B) and at the end (C) of Rhodamine B injection into pre-fabricated, vascular-like channels. (D) Experimental setup of the perfusion system in an incubator. Inset shows enlarged view of the perfusion chamber for liver model. Inlet and outlet of the channel network are connected to perfusion tubing through 18 gauge needles. Fluorescent images of live and dead cells in channelized, perfused liver model (E) and solid, statically-cultured liver model (F) after 3 days of culture. (G) Measured cell viability in channelized and solid liver models at both the center and edge regions, n = 10. (H) Albumin production for channelized liver model, solid liver model and casted model after 6 days of culture, n = 6. Cell-laden liver models in (E-G) were printed using 15% PEGDA 4k Da conjugated with 10mM RGD and liver model in H was printed using 8% GelMA plus 5% PEGDA 8K Da. All box plots with whiskers represent the data distribution based on five number summary (maximum, third quartile, median, first quartile, minimum). **, *p* < 0.001; *, *p* < 0.05; determined by non-parametric unpaired t-test.

To support the survival of cells encapsulated in the bulk hydrogel body of the liver model, the inlet and outlet of the channel network were connected to a recirculating flow system (Figure 4D). Media perfusion in the channel network was maintained for up to 6 days and cell viability and albumin secretion by encapsulated HepG2 cells were analyzed. As a comparison, a non-channelized liver model of identical size and scaffold composition was fabricated and maintained in static culture. At the end of culture period, both liver models were sliced horizontally and vertically to allow the examination of cell viability throughout the bulk hydrogel body. Gross observation showed uniform cell distribution in the liver model without obvious clumps or cavities, confirming the good cell deposition ability of the FLOAT process (Figure 4E, F). Results of live/dead assay showed that perfused model has relatively high cell viability of approximately 80% in both the core and the edge regions (Figure 4G); however, the cell viability in the core region of the non-channelized model is only 50%. The cell viability in the edge region of the non-channelized model is reasonably high at approximately 70%, possibly due to the direct contact with the culture media (Figure 4G). Albumin secretion of the perfused liver model was 5.2 μg per million cells which is nearly three times higher than that of the non-perfused liver model (Figure 4H). The casted hydrogel thin layer model was observed to secrete a slightly higher amount of albumin (6.5 μg per million cells) than the perfused liver model, possibly due to its small thickness (1 mm) that does not limit media diffusion. Together, these results suggested that the viability and physiological functions of cells encapsulated in FLOAT-printed, large-sized hydrogel model were maintained at reasonably high level under efficient media perfusion.

### Endothelialize the vessel network in FLOAT-printed hydrogel model

Since vascularization is critical for the long-term survival of engineered tissues in vivo (28), we explored the potential to endothelialize the pre-fabricated channels in FLOAT-printed thick hydrogel models. In an initial trial, HUVECs were found to adhere poorly on the channel wall in PEGDA models even with the pre-coating of fibronectin. To improve endothelial cell adhesion and spreading, we blended GelMA with PEGDA prepolymer and found that increased GelMA component along with reduced PEGDA component helped to improve endothelial adhesion. Using 7% GelMA blended with 3% PEGDA 8000 Da, we printed a tissue patch model (2 cm x 1 cm x 4 mm) consisting of two channels for endothelialization demonstration (Figure 5A). After seeding, a large amount of endothelial cells adhered to the channel walls and formed patches of aggregates that smoothened out in the first two days (Figure S10). A uniform and confluent layer of endothelial cells formed by day 3 and remained stable up to day 9 (Figure 5B; Figure S10). The cross-sectional view of the channels showed that endothelial cells fully lined the lumen of the microchannels (Figure 5C). The endothelial layer is of good quality at both the straight segment and junction areas of the channels with well-established endothelial cell junctions (Figure 5 D-F).

**Figure 5.**
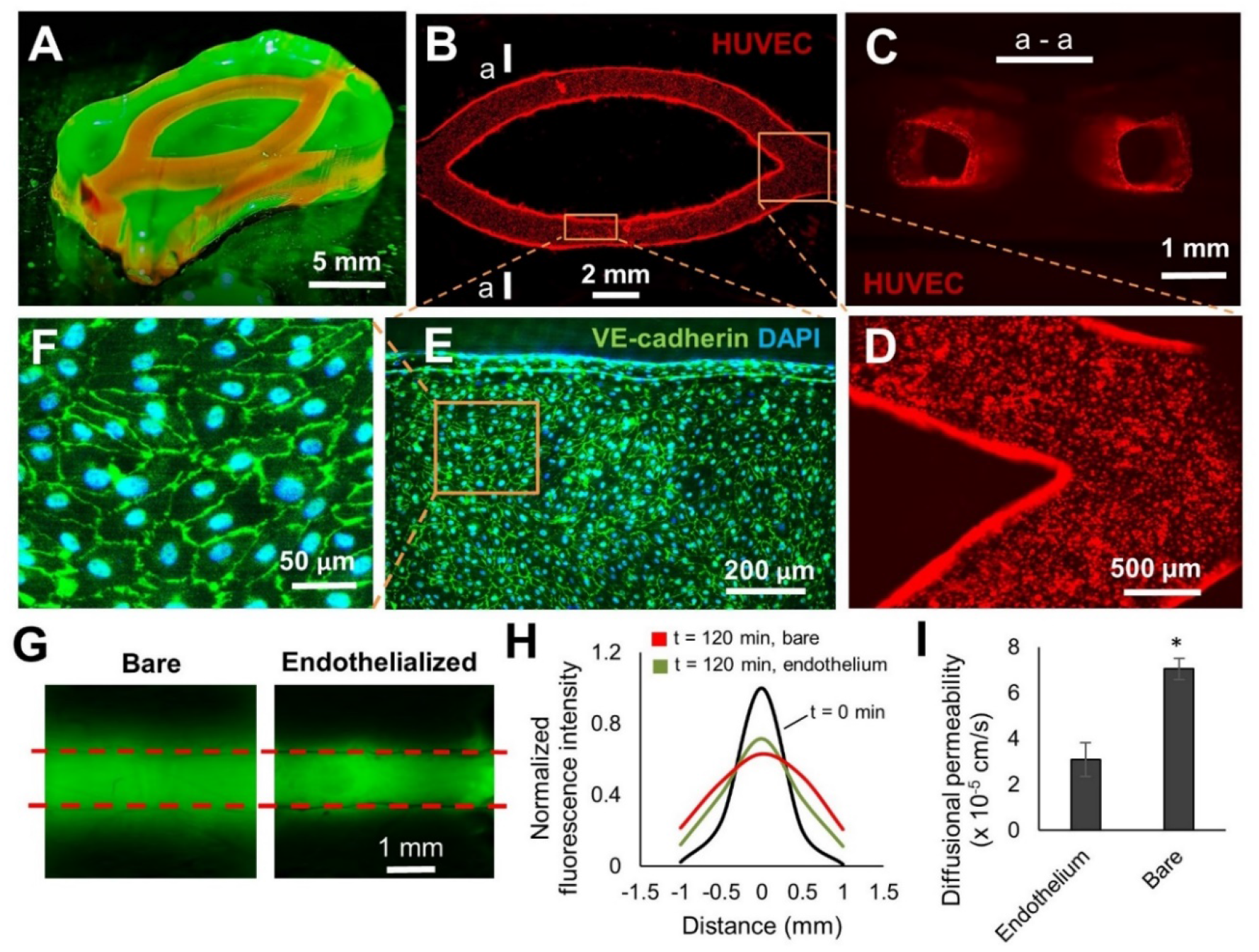
Endothelialize the printed vessel network in FLOAT-printed model. (A) A centimeter-size, channelized hydrogel model is illuminated under fluorescence. Channels are shown by injected Rhodamine dye. (B) A fluorescence image of HUVEC-lined channels 9 days after endothelial seeding. (C) The crosssectional view of endothelialized channels at section a – a shown in (B). (D) Zoom-in view of endothelialized corner region of the channels. (E) Immunofluorescent co-staining of the VE-cadherin and DAPI shows the formation of a confluent endothelial monolayer in the channel. (F) A zoom-in view of the cell layer shows well-formed endothelial cell junctions. (G) Fluorescence images show the diffusion of FITC-Dextran (3 – 5 kDa) in bare and endothelialized channels 120 mins after injection. (H) The plot of normalized fluorescence intensity change due to Dextran dye diffusion in the vicinity of a bare and an endothelialized channel over a 120 min period. (I) Calculated diffusional permeability of bare and endothelialized channels. Models in this figure were printed using 7% GelMA + 3% PEGDA 8 kDa. Data are reported as mean ± SD, n = 3, * p < 0.05.

Endothelial barrier function was assessed by measuring the rate of solute permeation from the lumen to the surrounding scaffold. The FITC-dextran (3-5kDa) in a bare channel was found to diffuse more rapidly and extensively than that in an endothelialized channel over a 120 min time period (Figure 5G, H). The diffusional permeability coefficient in bare channels was measured to be 7 x 10^-5^ cm·s^-1^, and this number decreased to 3 x 10^-5^ cm·s^-1^ in the presence of an endothelium (Figure 5I), which confirms the barrier function of the endothelium.

## Discussion

Hydrogel fabrication has been one of the main focused areas in bioprinting, but much of the efforts have been made on improving the printing resolution (28, 29). Although the spatial resolution limited by nozzle size and material morphology control has been improved recently for extrusion-printing, which allows the creation of tissue models with biomimetic features such as vascular channels (11, 14), such point-by-point material deposition method is still limited by its inability to print at multiple length scales and its slow printing speed that exposes cells to prolonged environmental stresses. The addition of colored dyes as photoabsorbers has allowed the fabrication of multivascular network using the layer-by-layer SLA method; however, this method still suffers from the slow printing speed, which requires anti-settling agents such as Xanthan gum and glycerol to prevent cell settlement during the fabrication of large-sized cell-laden tissue models (20). To address these challenges, FLOAT method combines high printing speed with multiscale printing capacity to allow the fabrication of a centimeter-sized hydrogel model in several minutes, as compared to several hours needed in extrusion-based printing and layer-by-layer SLA printing for a similar-sized part (11, 31). The development of the FLOAT method is achieved through studying the interactive effect of the process parameters and precisely controlling the prepolymer formula to enable high-velocity flow. Our studies demonstrated the different photopolymerization conditions that permit and inhibit the high-velocity flow, which allowed the formulation of the optimal prepolymer composition. The low suction force occurred in the optimal curing condition was shown to be critical to drive the prepolymer flow and to maintain good printing quality, thus unveiling the fundamental mechanism of the process.

In the current study, we showed that the continuous SLA process is unique to the hydrogel prepolymer solutions and cannot be achieved in resins without externally supplemented oxygen. It is possible that continuous hydrogel printing can also be achieved with the O2 permeable window and external oxygen supply as described in the CLIP setup (21), though the experimental setup will be more complicated. The current studies on the prepolymer flow and suction forces under controlled photopolymerization conditions should provide guidelines for such studies. Suction force is one of the limiting factors for the continuous printing of large-size models. We showed that the suction force in FLOAT hydrogel printing is several hundred times less than that in continuous resin printing. This can potentially be explained by the fluid mechanics theory of Stefan’s adhesion where separation force is inversely proportional to the cube of the separation distance and linearly proportional to the fluid viscosity (23, 32). Although we did not measure the dead zone thickness in our continuous resin printing, it should be smaller than that in the CLIP method (120 μm). Since the liquid flow layer thickness is more than 650 μm in the FLOAT method, the more than 5 times difference in the separation distance can lead to several hundred times difference in the separation force between the two methods. The difference in the viscosity (~10 cP for PEGDA MW 4000 prepolymer and 100 – 1000 cP for WI-resin) further contributes to the difference in the separation force. The low suction force in FLOAT method permits the fabrication of large, solid parts with good quality, thus improving over continuous resin printing where only small-size, thin-walled parts can be fabricated due to the large suction force of the WI-resin (21–23).

In the current study, we performed material optimization to seek hydrogel formula with both good printability and good biocompatibility. One of the major factors affecting the printability is the mechanical strength of the material. Sufficient mechanical strength is needed to avoid layer delamination or part breakage during continuous printing. Owing to the very low suction force in FLOAT, we show that a low, soft tissue-like mechanical strength (1 – 8 kPa) of the printed part is sufficient to support the continuous printing. In the current work, we added PEGDA to the GelMA mixture (total polymer less than 13% v/v) to enhance its mechanical strength. Future strategies to enhance the mechanical strength of polymeric materials may include improving photo-crosslinking chemistry (33) or developing nanomaterial doped polymer composites (34). However, it should be noted that high mechanical strength out of the stiffness range of the soft tissues (1 – 100 kPa) should be avoided for tissue engineering applications, since it will not support the growth and function of encapsulated cells. In studies that do not involve cell encapsulation, such as the nerve conduits fabricated by Zhu et al. (35), high stiffness hydrogel parts (300 kPa – 3000 kPa) have been produced by using high concentration precursor solutions (total polymer 32.5% v/v). Relatively high concentration precursor solutions (total polymer 20% v/v) was used to build cell-laden constructs using layer-by-layer SLA, but they only supported short term cell viability and function within 24 hours (20).

## Materials and Methods

### Printing system setup

The printing system presented in the current study is based on MP-SLA setup with significant modification to the hardware and software systems to enable a continuous printing process. The system utilized a bottom-up configuration where a digital light processor (DLP) device dynamically projects a digital mask image onto photo-sensitive material at the bottom of a transparent liquid tank. A high-resolution (1080P) dynamic micro-mirror device (DMD, Texas Instruments) was used as a dynamic digital mask generator, a blue LED (405 nm) or UV LED (385 nm) was used as the light source, and a high-speed FPGA chip and on-board RAM was used to store large volume of images and display the images with ultra-fast speed to enable reliable and smooth continuous printing. A holistic framework was developed with customized and integrated hardware, software and firmware to enable the ultra-fast and multiscale printing by leveraging the knowledge on hydrogel material property, process dynamics and light-matter interaction. In particular, dynamic synchronization of the image generation and the motion of build-platform were achieved using custom software, which allowed for high-speed continuous printing and eliminated layer-by-layer process. Grayscale mask image optimization and light calibration were conducted to achieve the highest accuracy and precision, which enables multi-scale printing. A custom-made tank of dimensions 6 cm x 4 cm x 2cm was fabricated using a transparent glass as the base. The glass was coated with polydimethylsiloxane (PDMS) to render the surface hydrophobic, thereby preventing attachment of the cured part to the bottom of the tank.

### Hydrogel material preparation

Polyethylene glycol diacrylate (PEGDA) MW 4000 Da and 8000 Da were synthesized following published method (36). Lithium phenyl-2,4,6-trimethylbenzoylphosphinate (LAP), a visible light photo-initiator, was synthesized following published method by Anseth group (37). To enhance cell adhesion onto PEGDA scaffold, cell adhesive Arginine-glycine-Aspartic acid-Serine (RGDS) peptide (Bachem) was conjugated to monoacrylated polyethylene glycol succinimidyl valerate (a-PEG-SVA, LaysanBio) following the method described by Turturro et al (38). Gelatin from porcine skin was purchased from Sigma. Gelatin methacrylate was synthesized using previously published protocol (39). The final product was dialyzed and freeze dried before use.

### Cell culture and maintenance

Human liver cell line HepG2/C3A (ATCC) was maintained in Minimum essential medium (MEM) supplemented with 10% FBS and 1% Penicillin/Streptomycin. RFP-tagged human umbilical vein endothelial cells (RFP-HUVECs, Angio-proteomie) were cultured in endothelial cell growth medium (EGM-2, Promocell). Human dermal fibroblasts were maintained using F12K media supplemented with 2% FBS. The human primary skeletal muscle cells were cultured in proliferation medium composed of high glucose Dulbecco’s Modified Eagle Medium (DMEM, Gibco) supplemented with 20% fetal bovine serum in a humidified 37 °C incubator at 5% CO2. hiPSC-MSCs were thawed in DMEM, high glucose, GlutaMAX™ Supplement, pyruvate media (Thermofisher, 10569010) with 10% FBS containing 10 μM ROCK inhibitor Y-27632 (MedChemExpress, HY-10583). After 24 hours, the cells were cultured in fresh media without the ROCK inhibitor. To obtain GFP-tagged HF-MSCs, lentivirus encoding GFP was produced using plasmid for GFP expression (CMV-GFP) purchased from Addgene (Cambridge MA). Transduction was carried out using CMV-GFP lentiviruses in growth media with 8 μg/ml polybrene (Sigma-Aldrich) for 3-4 hours.

### Cardiomyocyte Isolation

All animal experiments were performed according to the protocol approved by the University at Buffalo Institutional Animal Care and Use Committee. Primary mouse cardiomyocytes were harvested following manufacturer recommended procedure. Briefly, day 1-4 neonatal mice were euthanized by cervical dislocation and their hearts dissected. Cardiac tissue was minced into 1-3 mm^3^ pieces and digested using the Pierce Primary Cardiomyocyte Isolation Kit (ThermoScientific). For each experimental replicate, primary cardiomyocytes were pooled from 7 neonatal mouse hearts (yielding approximately 5-10×10^6^ cells). The entire isolation protocol was accomplished within two hours of animal euthanasia to ensure high cell yield and reproducible viability. Isolated neonatal mouse cardiomyocytes were cultured in DMEM for primary cell isolation (Thermofisher, 88287) supplemented with 10% heat inactivated FBS.

### Measurement of liquid flow velocity and uncured liquid layer thickness

The flow properties of the hydrogel prepolymer solution in the uncured liquid layer during FLOAT printing was studied by tracking the movement of the fluorescent microbeads in the solution. Green fluorescent microbeads (d ~ 25μm, Fluoresbrite, Polysciences) were mixed with 20% or 80% PEGDA prepolymer solution (MW 4000 Da) containing 0.1%, 0.6% and 1% photoinitiator LAP and 0.03% photoabsorber Orange G. Both 20% PEGDA and 80% PEGDA solutions were used, and two printing speeds of 50 μm/s and 125 μm/s were tested. The video of the microbead movement during continuous printing of a 4 mm wide part was recorded through a side-mounted inspection microscope at 5 frames per second. The bead flow trajectories were tracked using ImageJ, and the flow velocity was determined as the travel distance divided by the travel time. The uncured liquid layer thickness was defined as the distance between the bottom surface of the cured part and the glass tank base and was measured in the video. The bead flow velocity at different heights in uncured liquid layer was measured and compared.

### Measurement of suction force at the curing interface

We measured the suction forces at the curing interface during FLOAT printing of 10% and 20% PEGDA 4000 Da hydrogel pre-polymer solutions containing 0.6% LAP photoinitiator using an UXcell load cell sensor with a maximum capacity of 100 g (model # a11112800ux0213). The suction force during MP-SLA printing of acrylate-based resin part (Makerjuice Standard, MakerJuice Labs) was measured using an UXcell load cell sensor with a maximum capacity of 2 kg (model # a14071900ux0072). The load cell weighing sensor was mounted as a cantilever beam structure, with one end fixed to the Z stage and the other end fixed to the build platform. The suction force is recorded at three different platform moving velocities: 0.05 mm/s, 0.2 mm/s and 0.5 mm/s. A solid cube model with 1 cm edge length was used in the printing of both the PEGDA hydrogel and resin materials.

### Curing depth study

Curing depth is the thickness of the layer that is cured by light irradiation at critical exposure, and it is significantly affected by the exposure energy and the light absorbing property of the photosensitive material. We varied both the exposure energy and the type and concentration of the photoabsorbers to study their effects on the curing depth during FLOAT printing of PEGDA hydrogel. The base prepolymer solution contains 20% PEGDA 4000 Da, 0.6% LAP photoinitiator and 0.01% TEMPO. Five different photoabsorbers including 0.1% Orange G (Sigma), 0.1%, 0.15% and 0.2% Quinoline yellow (Sigma), 0.1% Tinuvin Carboprotect (a gift from BASF), 0.1% 4-hydroxy benzophenone (Sigma) and 0.1% benzotriazole (Sigma) were mixed with the base solution separately. Exposure energy was calculated as the production of energy density and the exposure time and was varied between 8 to 28 mJ/cm^2^. A rectangular image (4.7*9.4 mm) was projected for various exposure times, and the thickness of the cured layer was measured using a Verasonics ultrasound system coupled with an L22-8 linear transducer array at the frequency of 15 MHz. Furthermore, we assessed the effect of curing depth control on the printing resolution by printing a turbine rotor model under various photoabsorber conditions. The base of the turbine model is 10 mm and the thickness of the individual blades is 0.65 mm. The base prepolymer solution is the same as the one used in curing depth study. We tested Quinoline yellow at 0.05%, 0.2% and 0.3% and Orange G and HMBS at 0.2%.

### Optimization of hydrogel materials for FLOAT printing

To understand the compatibility of FLOAT printing with various hydrogel materials, we prepared pure PEGDA 4000 Da prepolymer with varying concentrations at 10%, 15%, 20% and 30% and pure GelMA prepolymer with two concentrations at 10% and 15%. We also prepared blended prepolymer containing GelMA and PEGDA 8000 Da at following concentrations: 6% + 2%, 7% + 3%, 8% + 5%, 10% + 2%, 10% + 5%, and 11% + 4%. Results show that blended prepolymers with less than 2% PEGDA were too soft to print, as demonstrated by the peeling of parts during printing.

### Measure the effect of the printing speed on cellular metabolic activity and cytotoxicity

The effect of the printing speed on cellular metabolic activity and cytotoxicity was assessed on cell-laden hydrogel models printed using either FLOAT method or layer-by-layer SLA method. Cell-laden solid hydrogel blocks (3mm x 3mm x 2 cm) were printed using both methods. A colorimetric XTT assay (2,3-Bis-(2-Methoxy-4-nitro-5-sulfophenyl)-2H-tetrazolium-5-carboxanilide disodium salt, Biotium, 10060; 5-Methylphenazinium methyl sulfate (coupling agent), VWR, 200000-760) was performed to determine the metabolic activity of cells, and an absorbance LDH assay (Lactate dehydrogenase assay, CytoTox-ONE™ Homogeneous Membrane Integrity Assay kit from Promega, G7890) was used to determine the level of cytotoxicity caused by membrane compromisation, Three different cell types that are commonly used in tissue engineering including hiPSC-MSCs, neonatal mouse cardiomyocytes and primary human skeletal muscle cells (hSMCs) were used in the printing and analyses. A prepolymer solution containing 7% GelMA plus 2% PEGDA 8000 Da, 0.6% LAP and 0.015% QY was mixed with the cell suspension. A solid hydrogel part with dimensions of 3*3*20 mm was 3D printed using either layer-by-layer SLA method (2 hours printing time) or FLOAT method (6 minutes printing time). The part printed using FLOAT method was maintained in an incubator for nearly two hours to wait for the completion of the part printed using layer-by-layer method. The two parts were then analyzed at the same time. The parts were sliced and digested with 5 mg/mL collagenase for 30 minutes to collect the cells for metabolic activity measurement using XTT assay. The supernatant was used to determine LDH activity.

### Centimeter-size liver model fabrication

The 3D computer model of the liver containing channel network was created in Rhinoceros 3D software and sliced. The channel network was designed based on the distribution of the main hepatic blood vessels but with modifications to facilitate media perfusion and nutrient delivery. The cross-section of a single channel is rectangular with a width of 0.8 mm – 1mm and a height of 3 mm. To print the liver model of 3.5 cm x 2.5 cm x 1.5 cm, 15 ml of 15% PEGDA 4000 Da prepolymer containing 0.6% LAP photoinitiator and 0.15% – 0.2% Quinoline yellow was prepared, with the supplement of 10 mM a-PEG-RGDS for cell adhesion. 8×10^6^ cells/mL of HepG2 cells were suspended in the above solution. Cell-laden liver models with or without vasculature channels were printed in a laminar flow biosafety cabinet and transferred to a P-100 dish containing culture media for long-term culture in a CO_2_ incubator. Both channelized and solid liver models were submerged in 10mL of culture media in the P-100 dish, but the channelized model was further supplied with media perfusion.

### Dye injection and diffusion in the liver model

To examine the interconnectivity of the channel network and the mass transport of small molecule solutes in the liver model, we injected Rhodamine B solution (1 mg/mL) through the inlet of the channel network using an 18-gauge needle. Dye diffusion over a period of 30 minutes was imaged under fluorescence at 5 min interval using a MVX10 MacroZoom microscope (Olympus) equipped with 1X Apo objective. Obvious diffusion of the Rhodamine into the bulk hydrogel scaffold was observed 5 minutes after injection. Time-dependent dye diffusion in the vicinity of a channel was quantified by measuring the fluorescence intensity.

### Recirculating flow system setup

During media perfusion of the channelized liver model, inlet and outlet of the vascular channels were connected to 1/16 in. sterile peristaltic tubings (PharMed BPT) through 18-gauge needles. Tubings were then hooked up to a Masterflex peristaltic pump (Cole Parmer). A 30ml syringe (Nordson EFD Inc.) was used in the fluidic circuit as a media reservoir. The total media volume in the fluidic circuit was about 20-25 ml. The entire fluid circuit operated inside a CO2 incubator at 37 degree under 0.4 mL/min flow rate to avoid disturbance to the construct. The channelized liver model was submerged in culture media in a P-100 dish during perfusion.

### Encapsulated cell viability and albumin production in cultured liver model

The relatively long-term cell viability in the liver models was assessed after 72 hours of culture using Live/Dead Viability Kit (L-3224, ThermoFisher). To ensure diffusion of staining dyes into the center of the centimeter-size liver model and also to better visualize the cells in the center region, models were sliced horizontally at the mid-height before staining. Live and dead cells were visualized on a fluorescence microscope, and the images from six representative areas at the edge and center of the liver models were used for quantification. The results were compared between channelized and solid liver models.

Albumin secretion by HepG2 cells encapsulated in liver models with or without vasculature channels was quantified using ELISA (Human Albumin ELISA kit, Ab 108788, Abcam). Liver models (polymer composition: 8% GelMA plus 5% PEGDA 8000 Da and 0.6% LAP) were FLOAT printed with a mixed cell population containing HepG2 cell to normal human dermal fibroblast at a ratio of 2:1 and a final density of 8 million cells per mL. The channelized model was maintained under media perfusion, and the solid model was maintained under static culture. Casted hydrogel thin layer of 5 mm x 5 mm x1 mm was used as a non-printed control. After six days in culture, the liver models were sliced through the horizontal and vertical axis, and sliced hydrogel parts were incubated for 3 hours to allow the release of the albumin from hydrogel core into the culture media. Cell culture supernatants were collected afterwards and used in ELISA assay. The absorbance of the samples was measured on a Sapphire microplate reader (TECAN) at 450 nm. Albumin standards were used to calculate the amount of albumin secreted by the cells. The concentration of albumin was then normalized based on HepG2 cell number.

### Endothelialization of vascular channel network

To demonstrate endothelialization of vascular-like channels, we FLOAT-printed a tissue patch model (2 cm x 1 cm x 4 mm) containing two microchannels, using 7% GelMA + 3% PEGDA 8000 Da mixed prepolymer solution composed of 0.6% LAP and 0.015% QY. Prior to HUVEC injection, 0.01 mg/ml fibronectin solution was injected into the channels and incubated for 45 min. A 50 μl suspension of RFP-HUVECs containing 10 x 10^6^ cells per mL was injected into the channels, and the cells were allowed to adhere for 5 hours and then placed on a shaker vibrating at 60 rpm for long-term culture in the incubator. The adhesion, growth and morphology of HUVECs in the channels were evaluated over a 9 day culture period.

### Diffusional permeability testing

We used FITC-conjugated dextran (MW 3 – 5 kDa) to study the diffusional permeability in the absence and presence of endothelial layer in the printed channels. EGM-2 media containing 100 μg/ml FITC dextran was injected into the channels with or without endothelial lining and the dye diffusion over a period of 2 hours was imaged under fluorescence at 30 min interval using a MVX10 MacroZoom microscope (Olympus) equipped with 1X Apo objective. The changes of the intensity profile over time were quantified using ImageJ.

Diffusional permeability, Pd (cms^-1^) of FITC-Dextran for a channel with diameter, d is quantified by using the equation (14, 40):

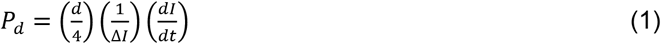

Where ΔI is the increase in the fluorescence intensity in the lumen right after injection of the dextran dye, dI/dt is the slope of the fluorescent intensity increase in the surrounding region with respect to time. Diffusional permeability of channels with and without endothelial lining were calculated and compared.

### Immunofluorescence staining and microscopy

Immunofluorescence staining was used to assess the uniformity of the endothelial layer in the vascular channels. After 9 days in culture, the printed models were fixed in 4% paraformaldehyde for 30 minutes and permeabilized with 0.1% Triton X-100 for 30 minutes at RT. The constructs were washed three times with PBS. Prior to incubation with primary antibody for VE-cadherin (ab33168, 1:500, Abcam), the constructs were placed in a blocking solution containing 3% BSA solution for 3 hours at RT. Primary antibodies to VE-cadherin was incubated overnight concentration in 1% BSA. The constructs were then washed with PBS three times to remove unbound primary antibodies followed by secondary blocking step with 10% Goat serum for 3 hours at RT. Alexa Fluor^®^ 488-conjugated anti-rabbit IgG (Abcam, ab150077, 1:400) was used as the secondary antibody. Samples were counterstained with DAPI to visualize nuclei. Confocal microscopy was performed on an Andor Technology DSD2 confocal unit coupled to an Olympus IX-81 motorized inverted microscope equipped with Plan-Apochromat ×10 air objective. Images were taken with an optical slice of 2 μm and 2D projected images were obtained using Z-stack tool on ImageJ.

### Magnetic resonance imaging (MRI)

MRI was performed on the printed hydrogel hand model using a 4.7 Tesla preclinical scanner incorporating the ParaVision 3.02 imaging platform and a 72 mm I.D. quadrature radiofrequency coil (Bruker BioSpin, Billerica MA). Prior to imaging, printed vascular channels were filled with mineral oil to aid in visualization of the channel morphology during MRI. Following scout scans, a three-dimensional, spoiled gradient echo scan was acquired encompassing the entire model. Acquisition parameters include: echo/repetition time = 3/15 ms, flip angle = 40 degrees, field of view = 64×50×40 mm, averages =2, acquisition matrix = 256×200×160.

Following acquisition, regions of interest were created for both the hand and channels using Analyze 10.0 (AnalyzeDirect, Overland Park, KS). A transparent, volumetric rendering was performed within Analyze using opacity values of 0.6 and 1.0 for the hand and channels, respectively.

### Statistics

Data are presented as the mean with error bars showing the standard deviation (SD). Non-parametric unpaired t-test or one-way ANOVA was used to determine statistical significance for two-group comparisons or multiple-group comparisons, respectively. Values of p<0.05 was considered to be statistically significant.

## Acknowledgments

Research reported in this study was supported by National Institute of Biomedical Imaging and Bioengineering of the National Institutes of Health under award number R01EB019411 (R.Z.). The content is solely the responsibility of the authors and does not necessarily represent the official views of the National Institutes of Health. Authors would also like to acknowledge the funding support from the School of Engineering and Jacobs School of Medicine and Biomedical Sciences at the University at Buffalo.

## Author Contributions

R.Z. and C.Z. conceived the idea, N.A., H.Y., Z.G., C.Z. and R.Z. designed the experiments, N.A., H.Y., Z.G., Z.C., K.W., J.L., N.R., S.A., Z.M., J.S., J.L., D.W., J.X., C.Z. and R.Z. performed experiments and analyzed data, N.A., H.Y., C.Z. and R.Z. wrote the manuscript. R.Z. and C.Z. supervised the project. N.A., H.Y., Z.G., contributed equally to this work.

## Data availability

The raw data required to reproduce the findings in this paper are available upon request to the corresponding author.

## Supplementary Information for

**Fig S1.**
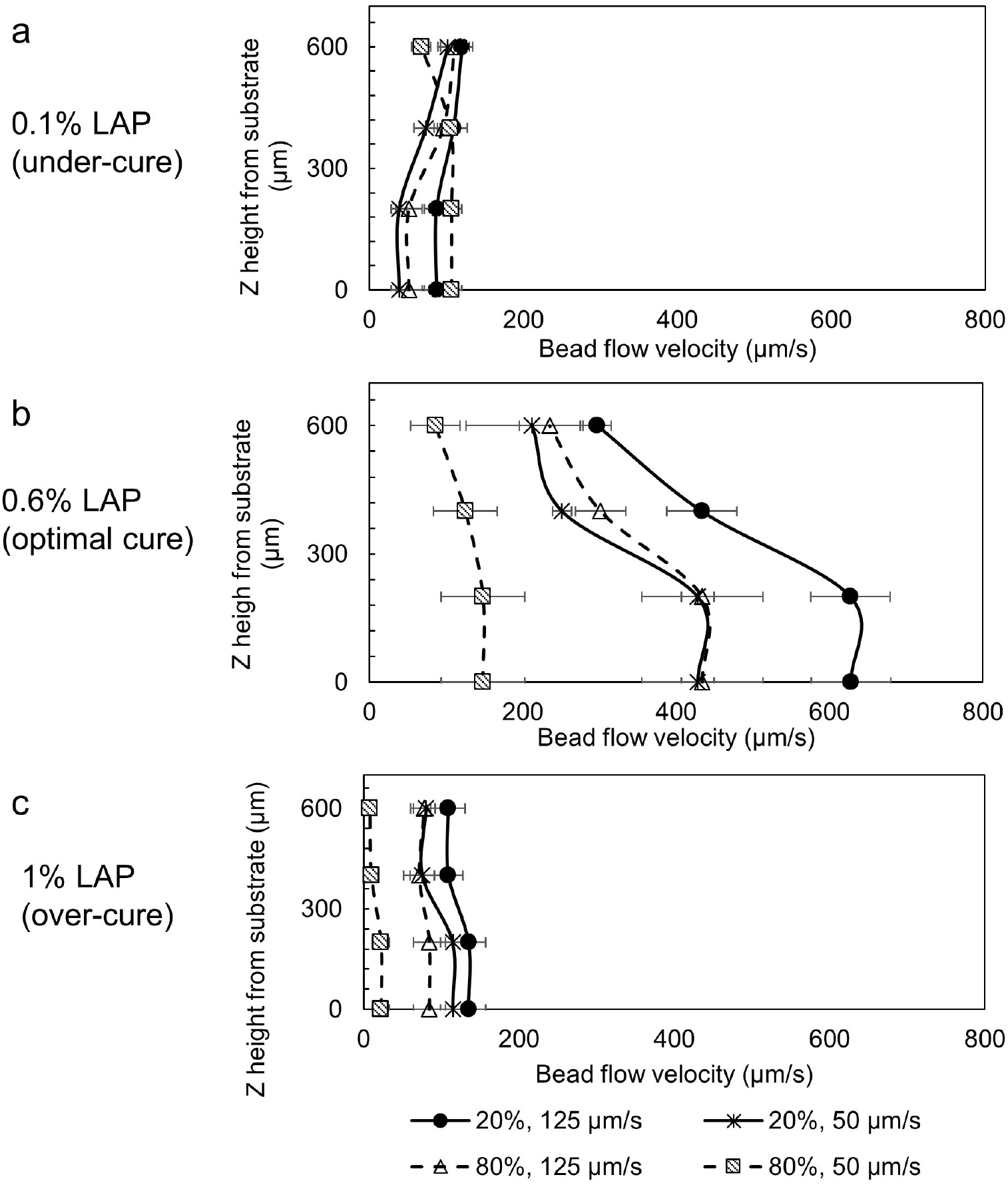
The flow velocity profile of PEGDA prepolymer solution under different photo-curing conditions. The measurement was performed by tracking the motion of the fluorescence microbeads in the photopolymerization zone just beneath the cured part. (A) Low photoinitiator concentration (0.1% LAP) results in undercured condition. The lack of photopolymerization greatly reduced the need for precursor material replenishment. As a result, only low-velocity, random and localized bead flow was observed. (B) In optimally cured condition (0.6% LAP), measured liquid flow velocity is in the order of several hundred micrometers per second and the highest flow velocity was observed in low concentration solution (20% PEGDA) under high printing speed (125 μm/s), while the lowest flow velocity was observed in high concentration solution (80% PEGDA) under low printing speed (50 μm/s). (C) High photoinitiator concentration (1% LAP) results in over-cured condition. In this case, fast photopolymerization results in the formation of a thin layer (1 – 2 mm thick) at the tank bottom that interferes with prepolymer flow. As a result, only low-velocity bead flow was observed.

**Fig S2.**
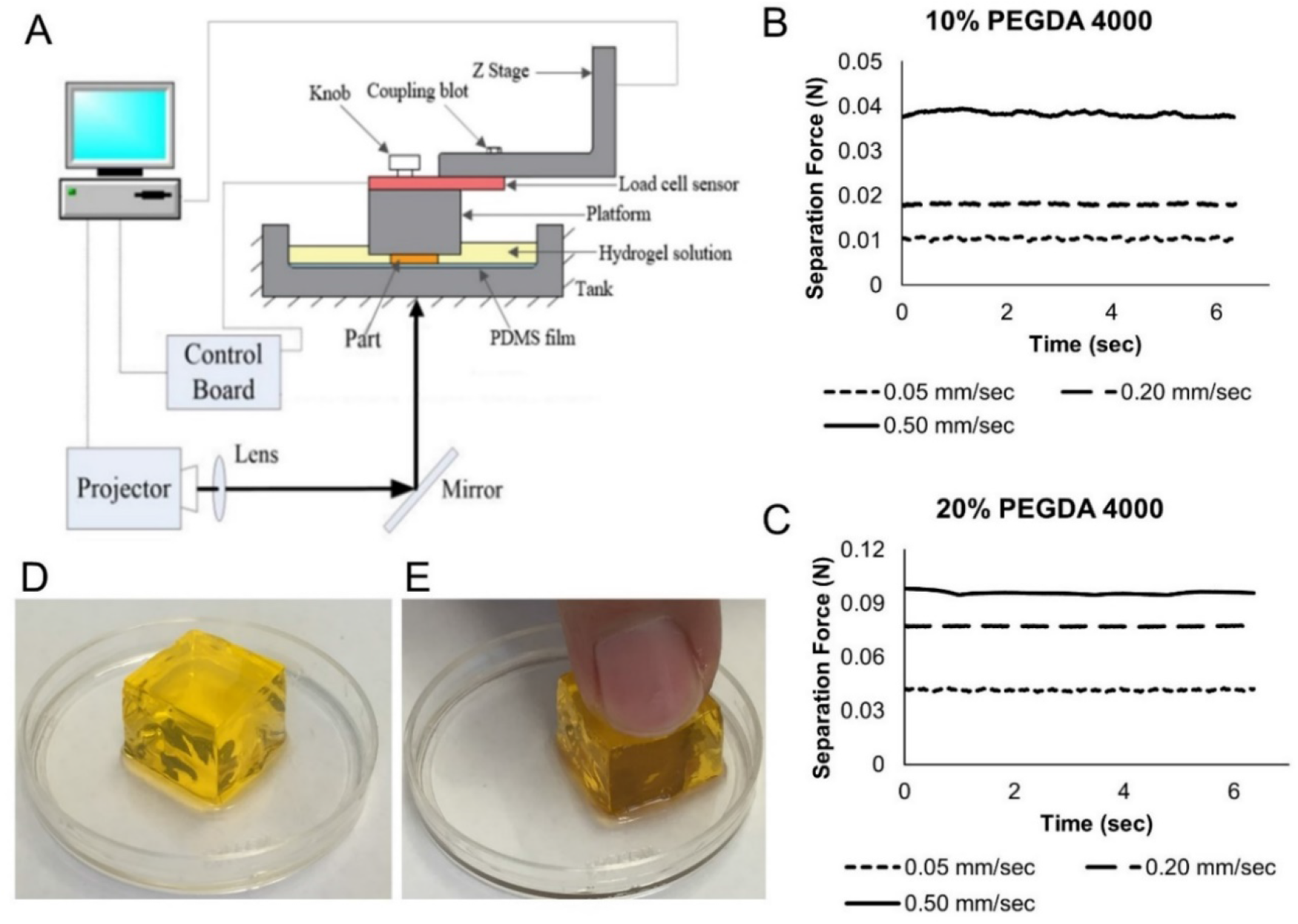
(A) Experimental setup for fluid suction force measurement. A force sensor was installed between the platform and the Z stage, and was used to report the suction force between the cured part and the tank base. Experimentally measured suction forces during FLOAT printing of 10% PEGDA (B) and 20% PEGDA (C) under different printing speeds. Note the suction forces maintain at a constant value during the FLOAT printing, indicating a stable printing process. (D) FLOAT-printed PEGDA cube of 1 cm edge length. Note the smooth surface and sharp edges of the hydrogel cube as a result of the stable printing process of the FLOAT. (E) PEGDA cube is very soft and can be easily deformed by finger compression.

**Fig S3.**
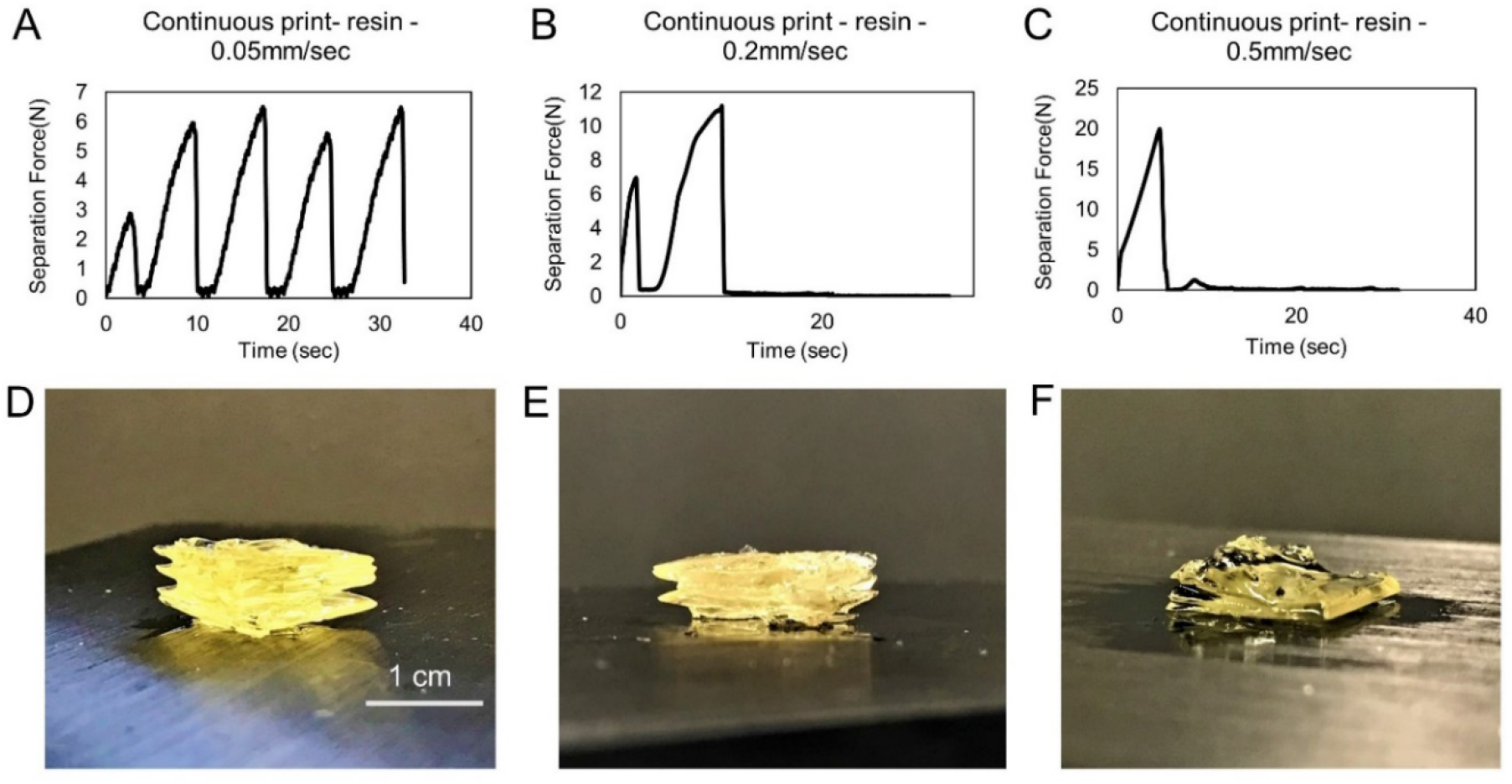
(A-C) Experimentally measured suction forces during continuous SLA printing of acrylic resin at various printing speeds. (D-F) Images of corresponding resin parts printed during suction force measurement. In panel (A), the highly wavy curve of the suction force at low printing speed (0.05 mm/s) indicates an unstable printing process which results in obvious delamination in the printed part (D). Note the number of peaks (4) in the suction force curve roughly matches the number of delaminated layers (3–4) in the part, suggesting the peak force occurs when the layer is attached to substrate and zero force (trough) occurs when the layer suddenly detaches from the substrate. In panels (B) and (C), only 1-2 force peaks were recorded, which correspond to 1-2 layers in the printed parts (E, F). The premature failure of the parts under these higher printing speeds (0.2 and 0.5 mm/s) is due to the very high suction forces that cause damage to the PDMS coating on the glass substrate.

**Fig S4.**
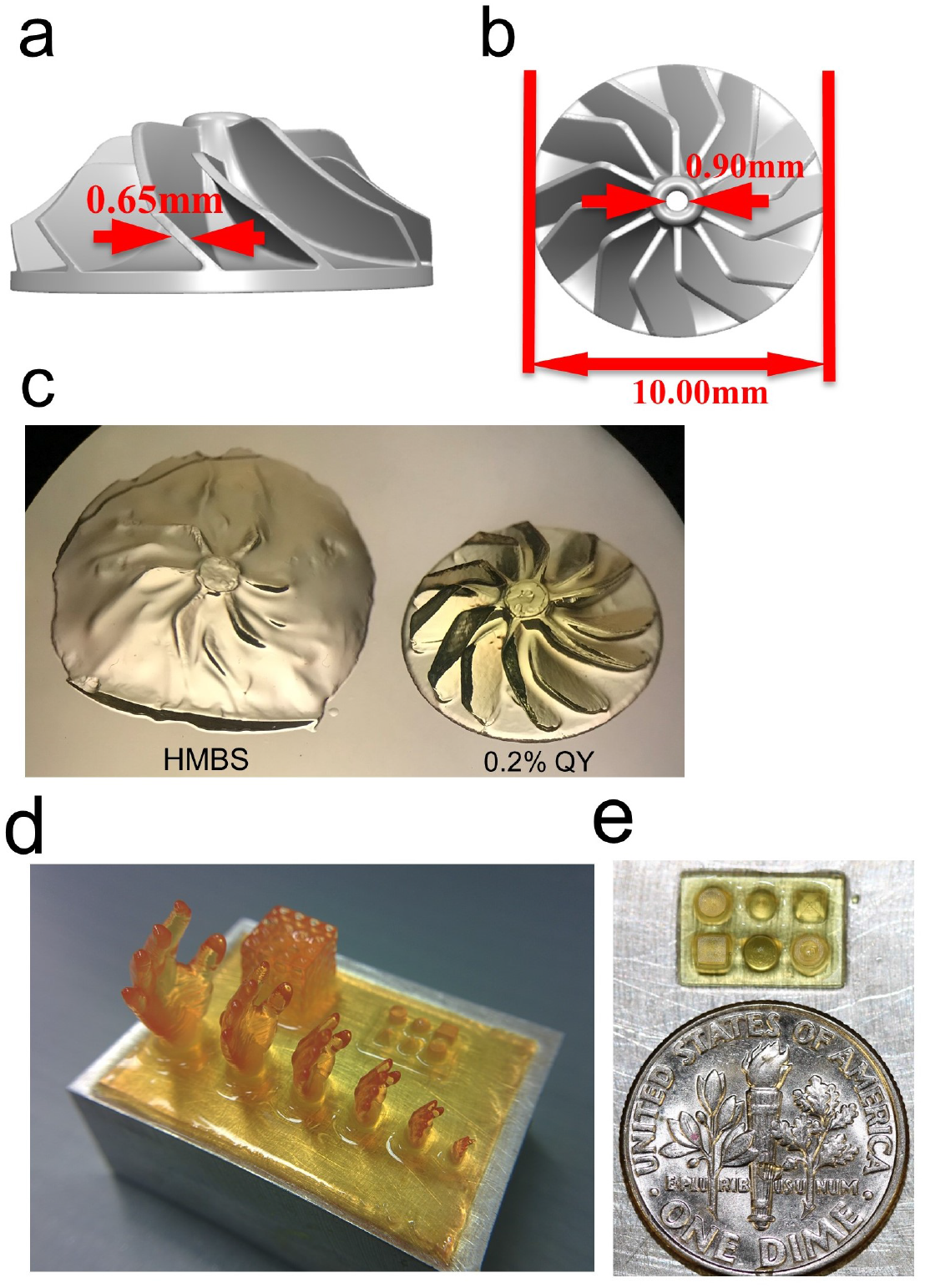
(A-B) Sideview and topview of the digital turbine rotor model with dimensions. (C) Pictures of printed turbine rotor models using optimal photoabsorber setting (0.2% QY) and HMBS. (D) FLOAT-printed PEGDA human hand models, a truss model and primitives viewed from hand model side. The base slab is 3.5 cm x 2.5 cm. (E) Zoom-in view of the primitives shown in panel A (From top left, cylinder, cone, pyramid, cube, dome, hollow cylinder). A U.S. dime coin (17.91 mm diameter) is placed side-by-side for comparison.

**Fig S5.**
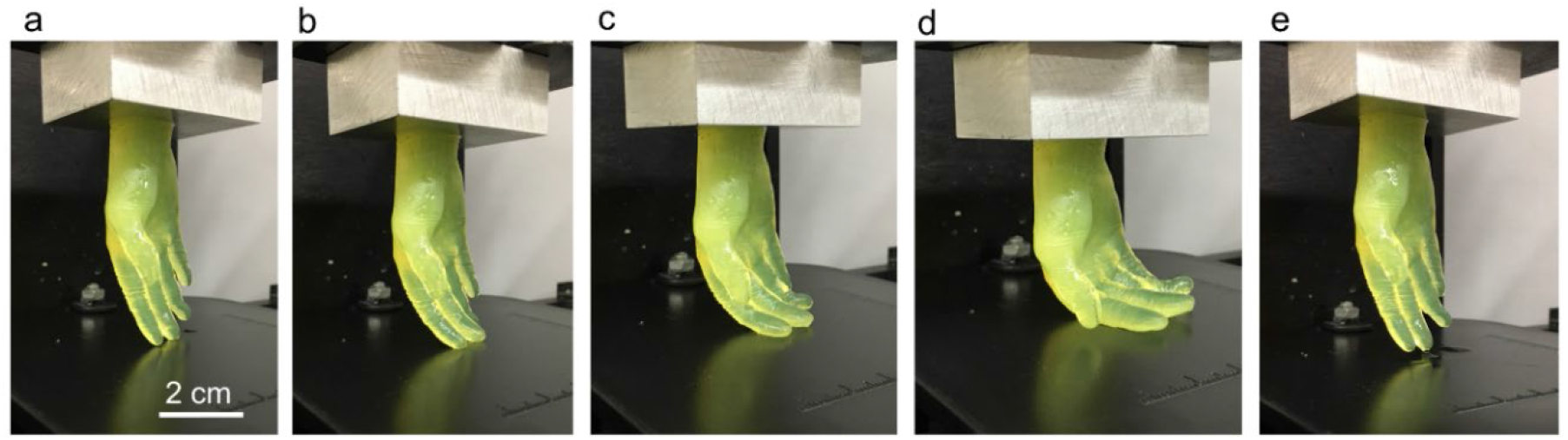
FLOAT-printed PEGDA human hand model under compression. (a-d) Sequential images show gradually increased bending of fingers under compression. (e) Recovery of the fingers upon compression release.

**Fig S6.**
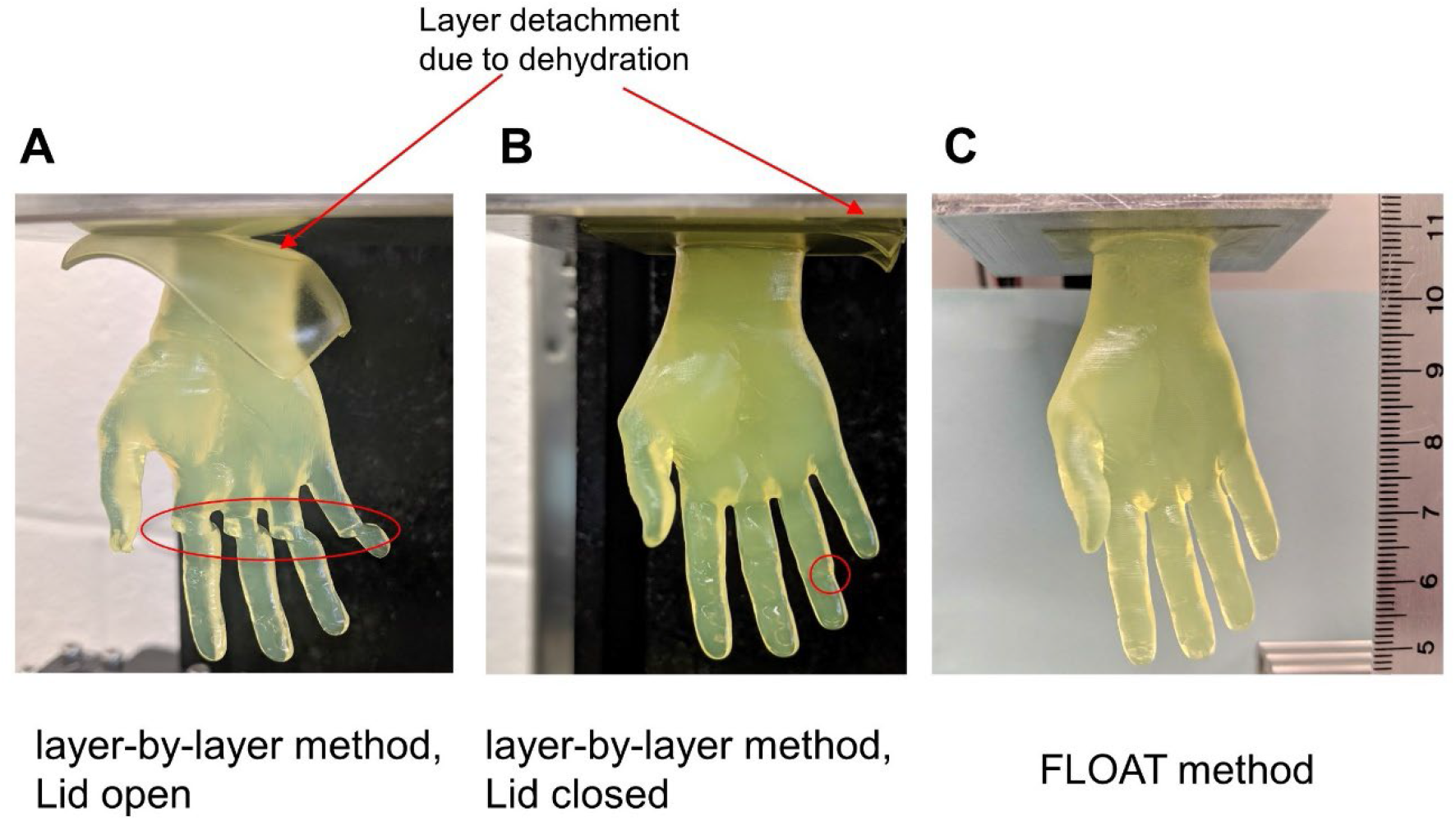
Comparison on the quality of a centimeter-sized hydrogel hand model printed using a traditional layer-by-layer based SLA method and FLOAT method. It took 2 hours to print the hand model using the traditional layer-by-layer based SLA process at 150 μm layer thickness. During this process, hydrogel dehydration occurred, causing layer detachment from the platform and misalignment in the fingers (red circles). (A) Hand model printed with printer lid open (rapid dehydration), severe layer detachment can be seen, which caused severe misalignment in the fingers. (B) Hand model printed with printer lid closed (slow dehydration). Mild layer detachment and finger misalignment can be seen. (C) Hand model is printed using FLOAT method in 19 mins. No dehydration is observed.

**Fig S7.**
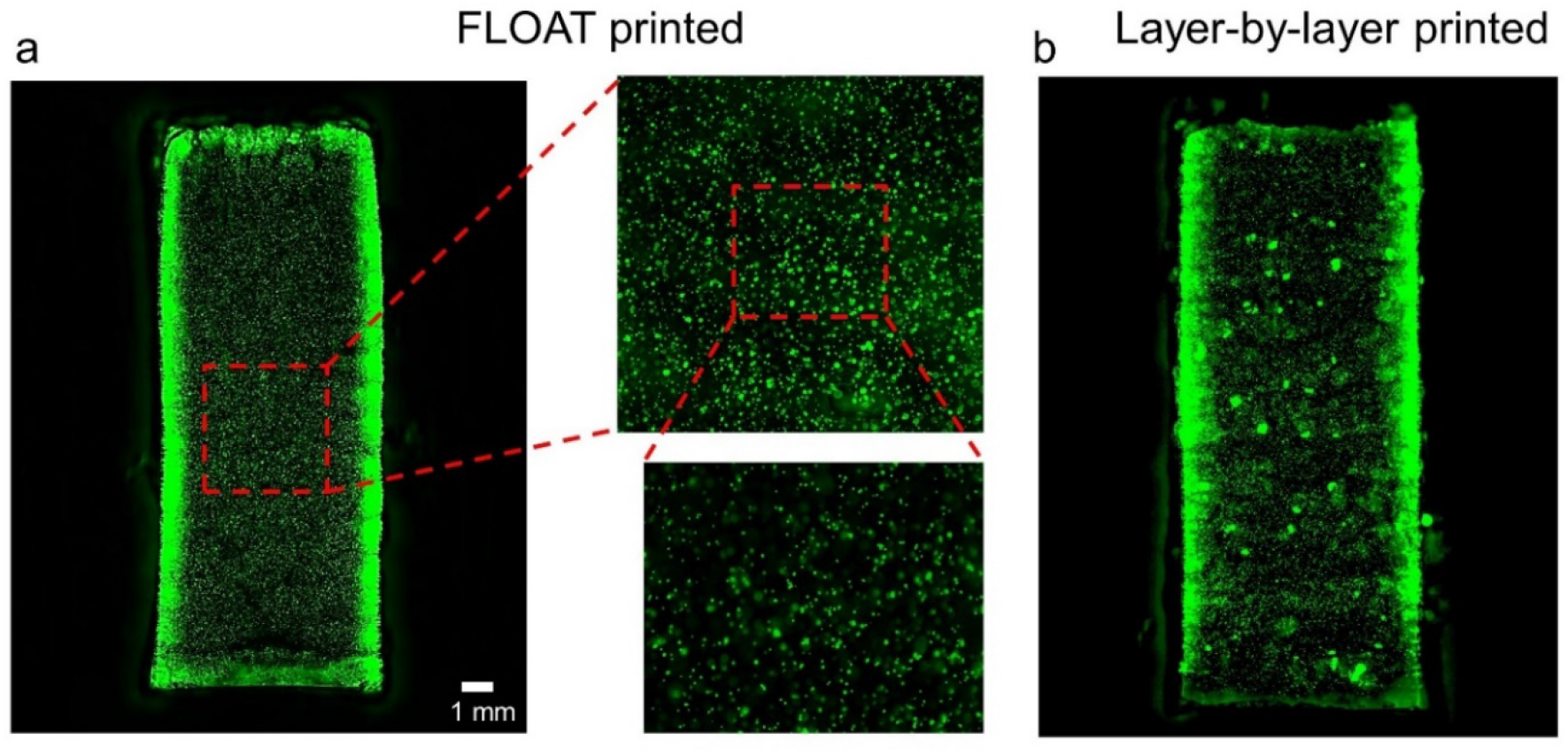
(a) Demonstration of the cell uniformity in FLOAT-printed centimeter-sized hydrogel part. The part size is 8*8*20 mm. The part was printed (platform moving direction) along its long axis. It took 6 mins to print this part. (b) Non-uniform cell distribution is seen in layer-by-layer printed part of the same size. Prepolymer solution containing cells was manually pipetted once in a while to prevent cell settling; however, layered cell distribution and cell aggregates were still seen in the part. It took 2 hours to print this part. Human skeletal muscle cells are used in both models.

**Fig S8.**
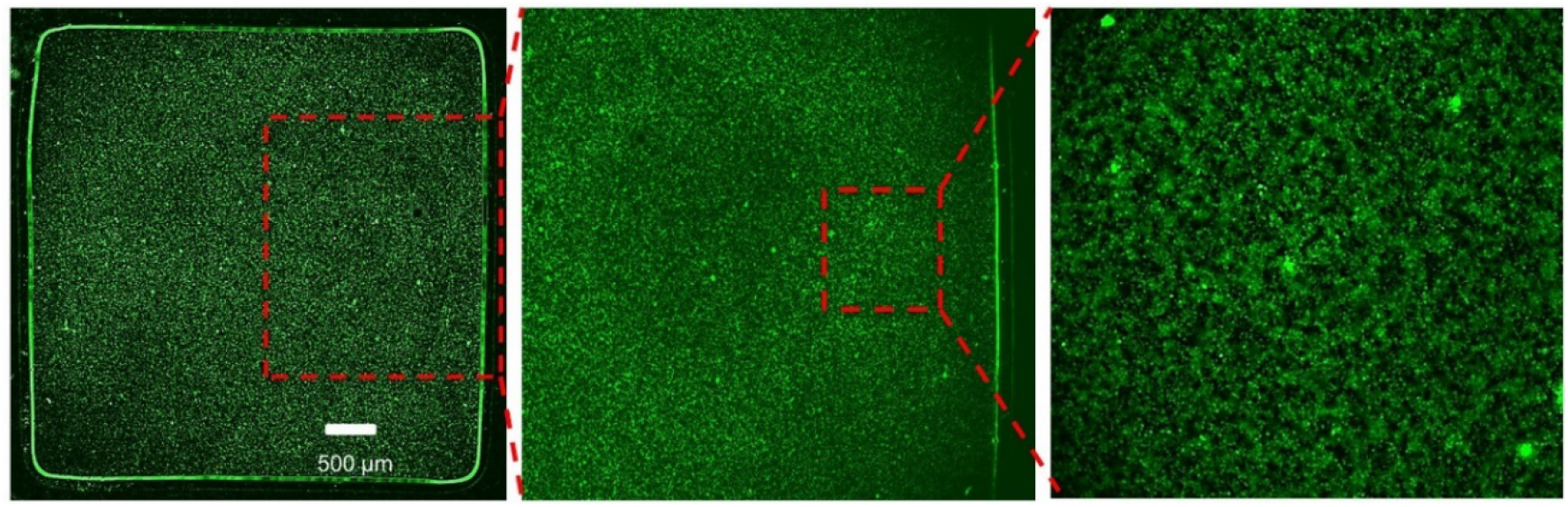
Demonstration of the ability of the FLOAT method to print at high cell density. Images show a centimeter-sized hydrogel part of 5*5*10 mm printed with 3T3 cells at a density of 50 million cells per mL.

**Fig S9.**
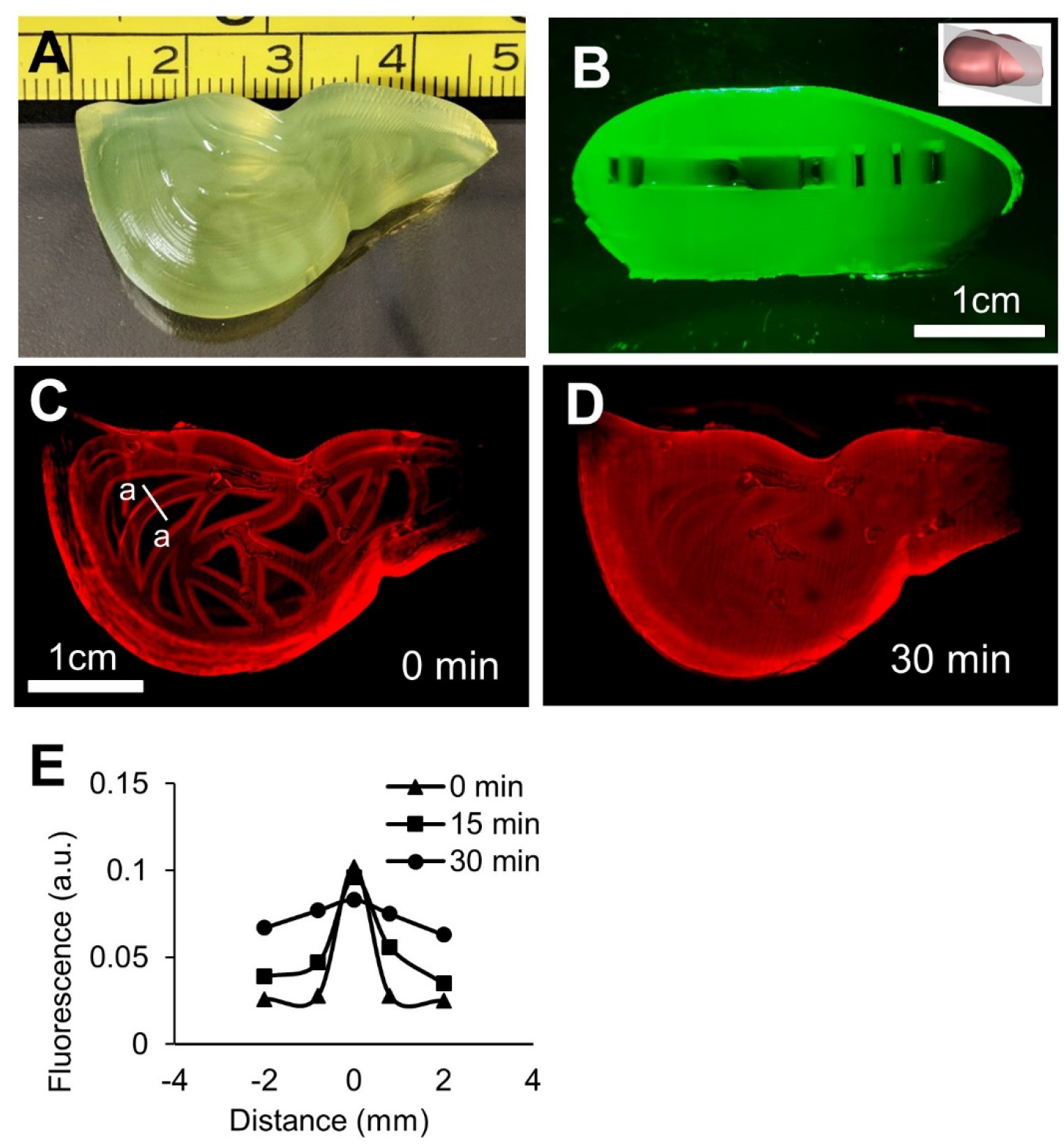
FLOAT printing of a centimeter-scale hydrogel liver model containing perfusable channels. (A) A liver model with smooth surface and monolithic, translucent hydrogel body was printed using 15% PEGDA 4 kDa. Measuring tape unit is in centimeter. (B) Cross-sectional view of the liver model to show the channel openings. Inset shows the direction of the cut. (C) Beginning of Rhodamine B dye diffusion. (D) Full diffusion of the dye into the interstitial space of the liver model was achieved 30 minutes after injection. (E) A representative plot of the fluorescence intensity change due to dye diffusion in the vicinity of a vascular channel (a-a cross-section shown in panel C) over a 30 minute period.

**Fig S10.**
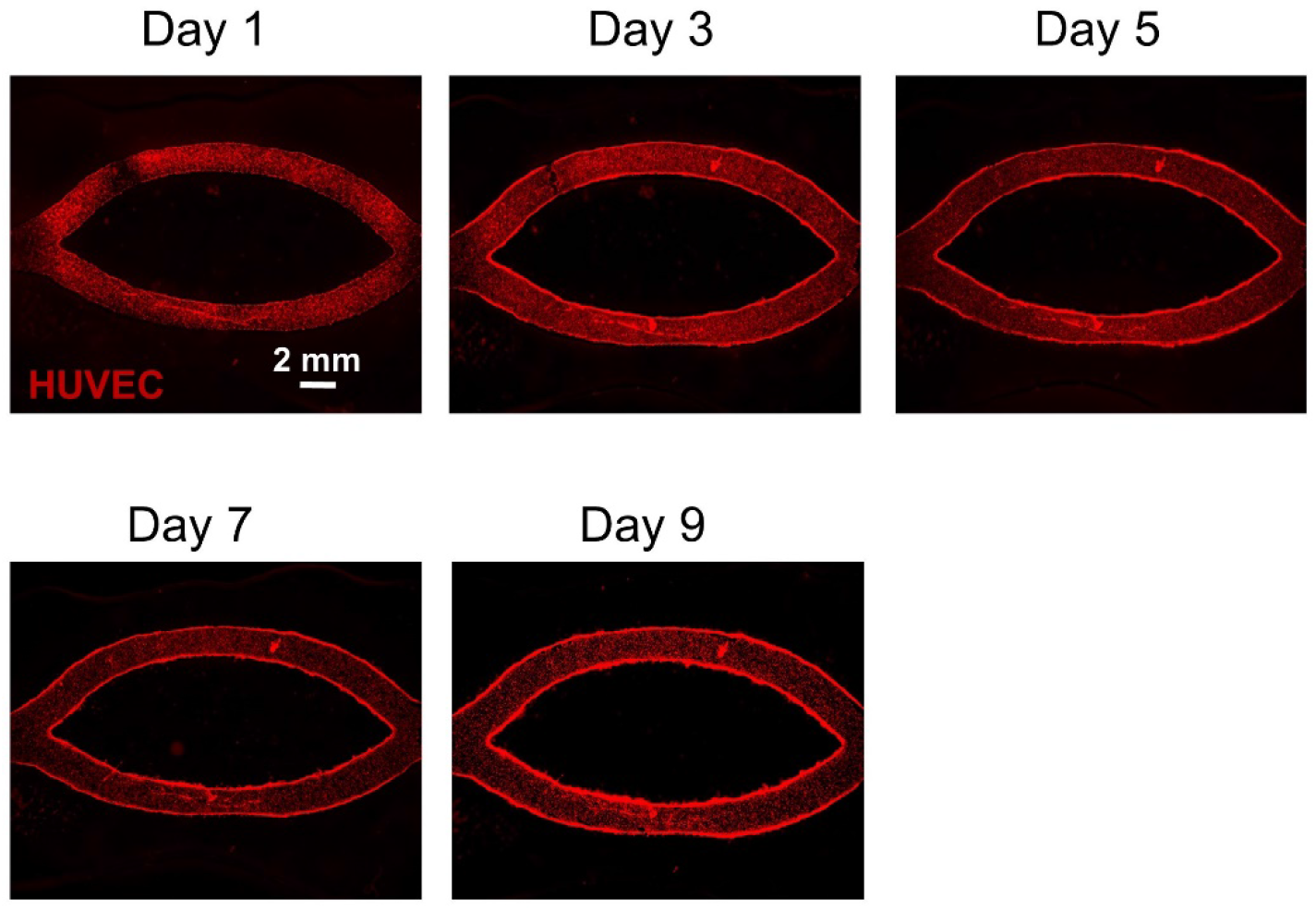
Time-lapsed images show the HUVEC coverage in printed channels in a hydrogel part. In the first three days after seeding, HUVEC coverage was found to be uniform in the channels except for a couple of isolated gaps. A uniform and confluent layer of endothelial cells formed after day 3 and remained stable up to day 9. This hydrogel model containing channels was printed using 7% GelMA + 3% PEGDA 8000.

## Notes

### Competing Interest Statement

The authors have declared no competing interest.

